# Single-cell sequencing of rodent ventral pallidum reveals diverse neuronal subtypes with non-canonical interregional continuity

**DOI:** 10.1101/2024.03.18.585611

**Authors:** David J. Ottenheimer, Rhiana C. Simon, Cassidy T. Burke, Anna J. Bowen, Susan M. Ferguson, Garret D. Stuber

## Abstract

The ventral pallidum (VP) was defined as a basal ganglia nucleus with dense input from ventral striatum. To further investigate a VP regional identity, we conducted a cross-species transcriptional characterization of VP cell types. We performed single nucleus RNA-sequencing of VP tissue from mice and rats and identified 16 VP neuronal subclasses with striking cross-species conservation. VP GABAergic neurons were surprisingly heterogeneous, consisting of 14 sub-classes from 3 developmental classes. Combining our sequencing data with a spatial atlas revealed that all VP subclasses extended beyond the traditional borders of VP. Integrating our VP data with prior sequencing data from striatal, hypothalamic, and extended amygdalar tissue confirmed that cell types are shared among these regions. Due to the role of VP in feeding behavior, we also assessed the transcriptional impact of high-fat diet consumption, which induced altered expression of genes involved in oxidative phosphorylation and inhibitory signaling. Overall, our results demonstrate that VP is not a transcriptionally discrete nucleus; rather, VP contains cell types with diverse expression patterns that overlap with regions beyond the basal ganglia.

## Introduction

The ventral pallidum (VP) was first identified as one of the major targets of nucleus accumbens (NAc) efferents (*1; 2*). This projection is a key node in the ventral striatopallidal system within the basal ganglia (*3–5*). Functional experiments have established the importance of this projection from NAc to VP for reward-related behaviors (*6; 7*), but, importantly, many studies have additionally demonstrated bidirectionality in this circuit (*8; 9*) and signaling in VP that appears independent of NAc (*10–12*), as well as important interactions between VP and other brain regions beyond the striatopallidal system (*13–20*). These observations, combined with a large body of work establishing VP as an important contributor to motivation, feeding, and reward processing (*21–23*), emphasize the importance of a thorough characterization of VP to better define its characteristics as an independent brain region.

A major challenge for VP research is delineating its borders. Owing to its origins as the output of NAc, VP is traditionally identified by the presence of Substance P, released by NAc fibers, and is otherwise defined by partial overlap with substantia innominata, extended amygdala, and basal forebrain magnocellular complex (*22*). Thus, an important question is whether VP contains cell types that are genetically distinct from nearby regions. Within VP, several markers have been used to define neural subpopulations, including markers for the neurotransmitters GABA, glutamate, and acetylcholine (*22*), and these accompany a collection of compelling findings on functional differences between these populations (*16–18; 24*). The genetic diversity within neurotransmitter cell classes is unknown, as is the distribution of other neuronal markers across these classes. Many studies have examined VP in mice and rats (*23*); however, little is known about the conservation of VP cell types between the two species.

To address these questions, we performed single nucleus RNA-sequencing of tissue from adult male and female mouse and rat VP. This approach allowed us to characterize the diversity of cell types in VP and the consistency across species. We integrated our results with the Allen Brain Cell Atlas (*25*) to standardize labeling of VP neuronal subclasses and assess their spatial distribution in the rodent brain. VP contained 16 neuronal subclasses, the majority of which were GABAergic. All these subclasses extended into adjacent regions, including NAc, the extended amygdalar bed nucleus of the stria terminalis (BNST), and the hypothalamic preoptic area (POA), a finding we validated by integrating our VP data with three additional RNA-sequencing datasets. Within VP, there was remarkable homology between mouse and rat tissue, both in subclass representation and marker gene expression, including developmental transcription factors and extracellular signaling proteins. Guided by the general role of VP in feeding (*21–23*) and specific alterations to VP following high-fat diet consumption (*26; 27*), we assessed the impact of prolonged exposure to high fat diet on VP transcription, and we found changes in the expression of genes related to oxidative phosphorylation and inhibitory signaling mechanisms. Our results demonstrate considerable transcriptional heterogeneity in VP as well as a gradient of subclass representation across space, eschewing anatomical boundaries, lending new insight into the regional identity of VP and its incomplete separation from adjacent (including non-basal ganglia) areas.

## Results

### Single nucleus RNA-sequencing reveals VP cell types

Our first goal was to identify populations of VP neurons with distinct gene expression patterns. To this end, we used single nucleus RNA-sequencing to measure cell-specific gene expression in VP. We dissected tissue from 24 mice and 4 rats (equal numbers of males and females), isolated nuclei, and prepared cDNA libraries using the 10x Chromium platform (Fig. 1A). Our dissections spanned most of the AP axis of VP (Fig. S1A). Due to the shape of VP, our dissections also included portions of ventral striatum (STRv) (Fig. S1B), but this permitted within-experiment comparisons of VP and STRv nuclei. Nuclei from mouse and rat tissue were of high quality, and canonical cell type markers allowed straightforward identification of neurons and glia (Fig. S2).

**Figure 1.**
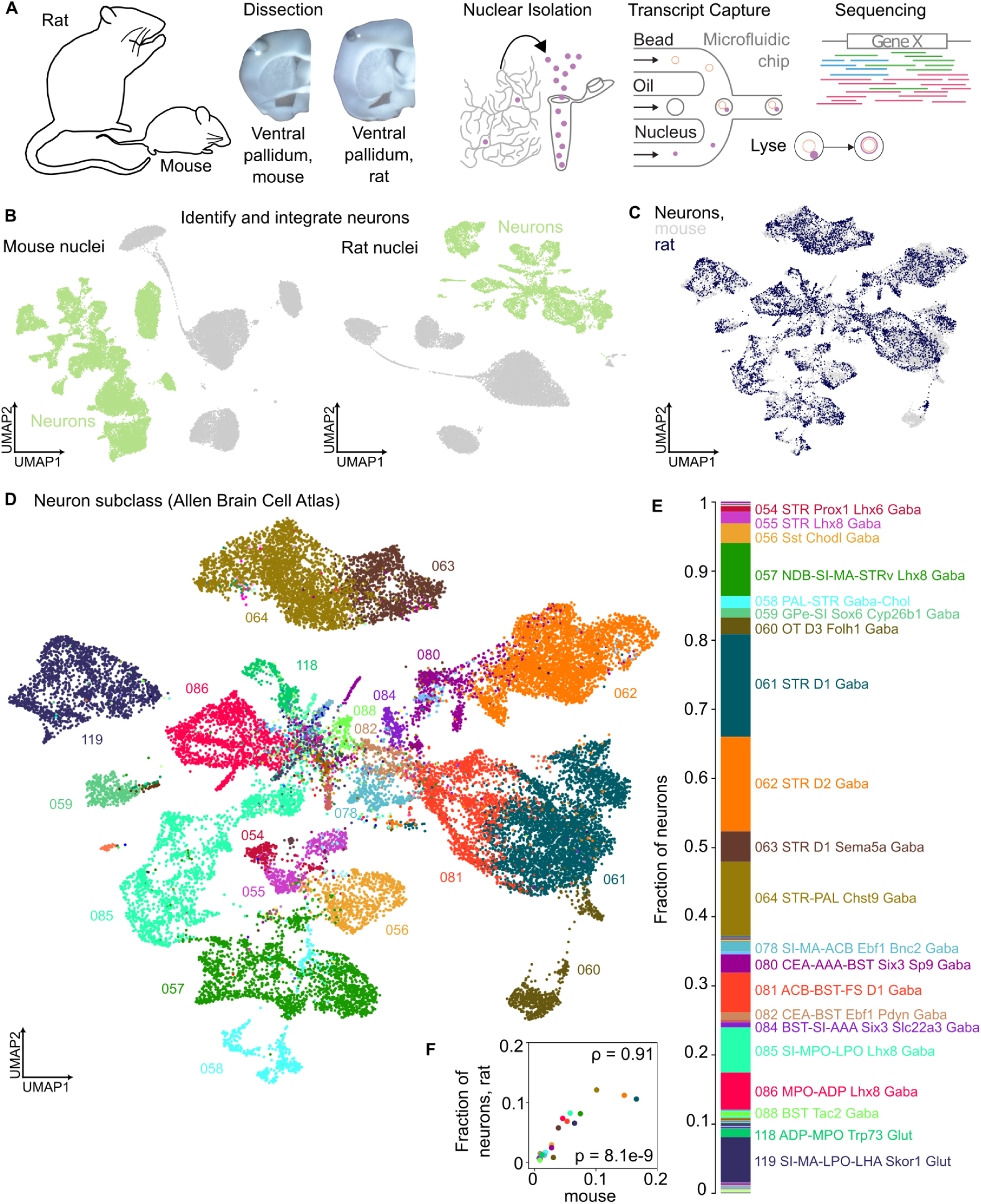
Single nucleus RNA-sequencing reveals VP cell types. (A) Process for sequencing mouse and rat VP nuclei. (B) All nuclei from mouse (left) and rat (right) VP dissections, plotted according to their gene expression similarity. (C) Neuronal nuclei from mouse and rat VP dissections, plotted according to their gene expression similarity and colored by species. (D) As in C, colored by Allen Brain Cell Atlas subclass. (E) The fraction of all VP dissection nuclei belonging to each subclass. Subclasses containing 0.5% of neurons are labeled. (F) The fraction of all VP dissection nuclei belonging to each subclass, separated for mouse and rat, with Spearman’s *ρ* and p-value denoted.

We next classified the neurons in our dataset into neuronal subtypes. First, we combined the identified neurons from mouse (*n* = 19, 588) and rat (*n* = 8, 143) into one dataset (Fig. 1B), containing 14,926 genes common to both species. Nuclei from both species spanned similar gene expression patterns (Fig. 1C). To standardize our results and maximize comparability with other sequencing studies, we integrated our data with the Allen Brain Cell (ABC) Atlas (*25*) using their MapMyCells tool. The subclass level of atlas classification was particularly useful, dividing neurons into many previously documented cell types, including striatal D1 and D2 medium spiny neurons and pallidal cholinergic neurons. Nuclei assigned to the same subclass clustered together in our dataset (Fig. 1D), indicating that similarity in gene expression patterns was preserved in the subclass assignments. In total, there were 21 subclasses each of which containing at least 0.5% of the neurons we sequenced (Fig. 1E). All 21 were present in both mouse and rat with similarly ranked proportions (Fig. 1F).

We validated our neuron classification by manually integrating our VP neurons with neurons from the ABC Atlas pallidum (PAL) dissections (*n* = 58, 337). We also included neurons from mouse NAc (*n* = 21, 474) (*28*) and our own dissections from rat NAc (*n* = 12, 312 neurons) (Fig. S3A) to further evaluate classification consistency across regions and rodent species. Neurons from the ABC Atlas spanned the full range of gene expression patterns in the other datasets (Fig. S3B), permitting the assignment of ABC Atlas designations to the remaining neurons (Fig. S3C-E). This approach yielded remarkably similar neuronal subclass classification as MapMyCells, evident in high correlation in subclass representation between methods and species but not between different regions (Fig. S3F-H), supporting the validity of using MapMyCells to effectively classify neurons from different regions and rodent species.

### A spatial gradient of VP neuronal subclasses

Examination of the spatial distribution of VP neuronal subclasses revealed a surprising lack of regional specificity. We leveraged the MERFISH-generated spatial component of the ABC Atlas to assess subclass location (Fig. 2A). First, we validated the presence of the 21 subclasses in VP and counted the number of neurons belonging to each subclass within the designated VP boundaries in the 4 MERFISH datasets containing VP. All 21 subclasses were detected in VP in the MERFISH datasets in similar proportions to our RNA-seq data (Figs. 2B-C). Furthermore, these 21 subclasses accounted for, on average, 95.9% of the neurons attributed to VP in the MERFISH datasets. Upon reviewing the spatial distribution of each subclass, we determined that 16 subclasses were in fact present in VP (Fig. 2D), while the remaining 5 were only present in striatum and were detected in our sequencing due to the adjacency of STRv in our dissections (Fig. S1B) and in the MERFISH counts due to imprecision in region boundary drawing (Fig. 2E). Overall, these results validated that we successfully sequenced and classified the primary VP neuronal subclasses.

**Figure 2.**
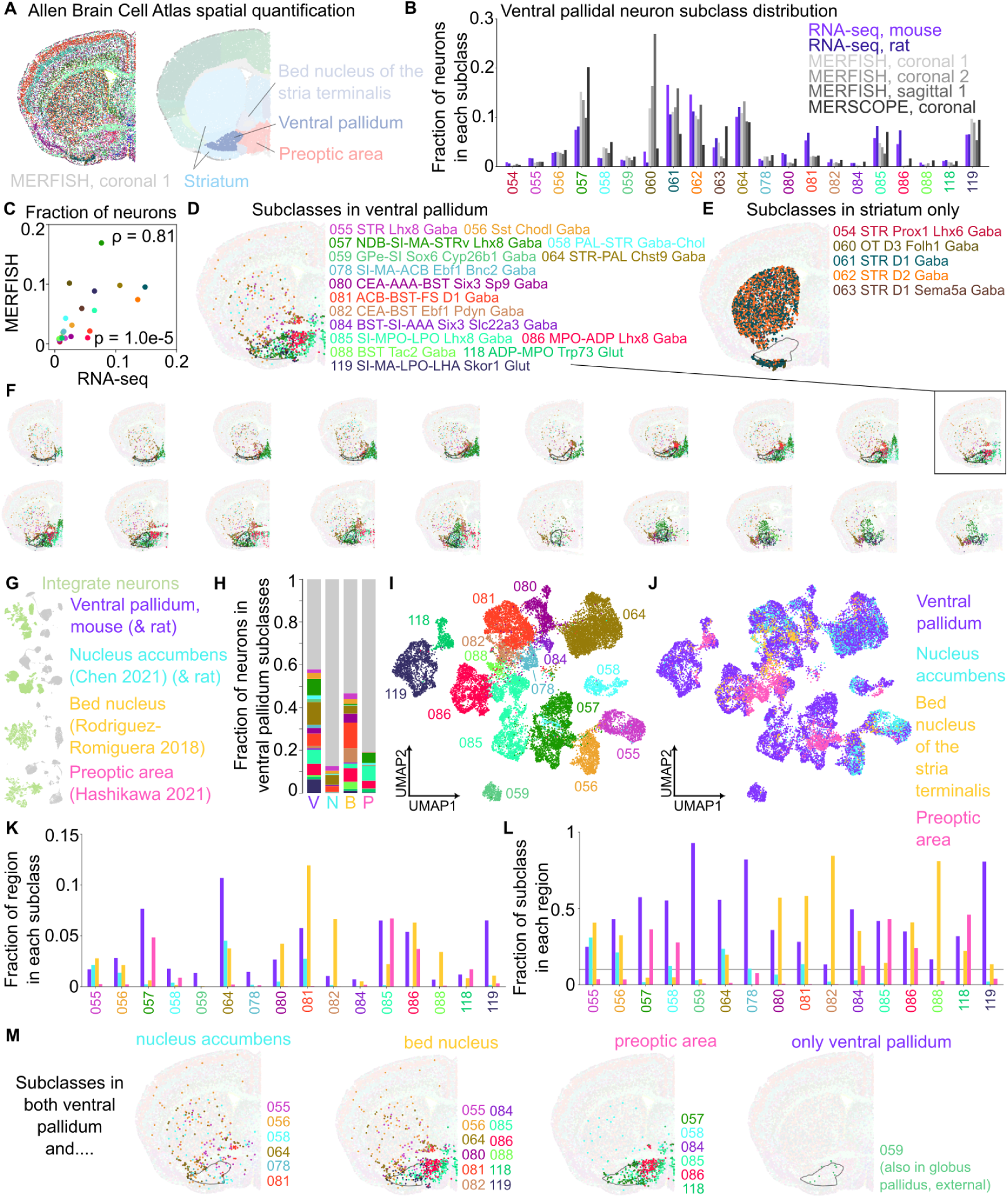
A spatial gradient of VP neuronal subclasses. (A) Example MERFISH section from the Allen Brain Cell (ABC) Atlas, colored by sub-class (left) and region (right). (B) The number of neurons belonging to each subclass we identified in Fig. 1 for our RNA-seq data from mouse and rat VP dissections and for the 4 MERFISH datasets containing VP (substantia innominata and magnocellular nucleus) in the ABC Atlas, as a fraction of all neurons in the dissection or region boundaries. (C) Fraction of all neurons in RNA-seq or MERFISH datasets belonging to each subclass, with Spearman’s *ρ* and p-value denoted. (D) Example ABC Atlas MERFISH section depicting all neurons belonging to VP sub-classes, with VP outlined in black. (E) Example ABC Atlas MERFISH section depicting all neurons belonging to the remaining subclasses, which were only present in striatum. (F) Neurons in VP subclasses across 20 consecutive ABC Atlas MERFISH sections containing VP (outlined in black). (G) UMAP plots of cells from the 4 mouse datasets we analyzed, with neurons highlighted in green. (H) Fraction of neurons from each dissection region (including rat VP and NAc) belonging to VP subclasses, assigned with MapMyCells. (I) UMAP plot of neurons from all dissection regions belonging to VP subclasses, colored by subclass. (J) As in (I), colored by dissection region. (K) Fraction of neurons from each region belonging to each VP subclass. (L) Fraction of neurons from each VP subclass belonging to each region (normalized by total neurons from each region). (M) Example ABC Atlas MERFISH sections depicting neurons belonging to VP sub-classes in NAc, BNST, POA, or none.

Further review of the 16 VP subclasses revealed diverse spatial distributions that extended beyond the traditional boundaries of VP into adjacent regions, including NAc, BNST, and POA (Fig. 2F). This observation introduced an unexplored possibility that VP contains neuronal populations that are transcriptionally continuous with nearby regions. We assessed this by integrating our VP neurons with neurons from NAc (as above), BNST (*n* = 1, 936) (*29*), and POA (*n* = 9, 807) (*30*) (Fig. 2G). We classified neurons from all 4 regions with MapMyCells and found substantial fractions of neurons belonging to our 16 VP subclasses in NAc (12.7%), BNST (46.7%), and POA (19.4%), compared to 57.9% in VP (Fig. 2H). The intermixed gene expression patterns across regions were visible when plotting all neurons from these subclasses (Figs. 2I-J). The proportion of neurons in each subclass differed across regions, however (Fig. 2K). Although VP neurons made up more than 10% of the neurons in all 16 subclasses, neurons from other regions were similarly represented in fewer subclasses: 6 in NAc, 12 in BNST, and 6 in POA (Fig. 2L). The spatial distribution of these subclasses in the MERFISH data confirmed the presence of each subclass in those regions (Fig. 2M). These distributions indicate that VP borders delineated by NAc axonal projection patterns do not correspond to a transcriptionally discrete brain region, although the precise representation and proportion of subclasses in VP is unique.

### Myriad genetic cell type markers among VP neuronal subclasses

By closely examining expression patterns in nuclei from our VP dissections, we uncovered remarkable heterogeneity in VP neuronal types. Guided by the subclass spatial patterns above, and knowing our VP dissections included STRv tissue, we assigned the neurons from our VP dissections to VP, STRv, both, or neither (Fig. 3A). Our ABC Atlas integration permitted simultaneous examination of neurotransmitter, ABC Atlas class, and ABC Atlas subclass in these populations (Figs. 3B-D). After correcting for the presence of STRv subclasses in our VP dissections, we were able to estimate the distribution of VP neurons across these categories (Figs. 3E-G). VP contained primarily GABAergic neurons (81.2%) but also glutamatergic (14.5%) and cholinergic neurons (2.4%), reflecting the known neurotransmitter distribution (*17; 23*), as well as a small population of GABA-glutamate co-releasing neurons (1.8%). The 14 GABAergic subclasses present in VP could be divided into 3 major classes corresponding to their development origin: 08 CNU-MGE GABA (23% of neurons, originating from medial ganglionic eminence, and also containing the cholinergic neurons), 09 CNU-LGE GABA (19.2%, lateral ganglionic eminence), and 11 CNU-HYa GABA (43%, anterior hypothalamic area, purportedly preoptic area in origin) (*25*). Over half of VP neurons were members of the 4 most numerous GABAergic subclasses: 057 NDB-SI-MA-STRv Lhx8 (14%), 064 STR-PAL Chst9 Gaba (16.7%), 085 SI-MPO-LPO Lhx8 Gaba (12.3%), and 086 MPO-ADP Lhx8 Gaba (10.2%). There were 2 glutamatergic subclasses, both members of the 13 CNU-HYa Glut class (14.5%), with 119 SI-MA-LPO-LHA Skor1 Glut containing the majority of glutamatergic neurons (12.3% of all VP neurons).

**Figure 3.**
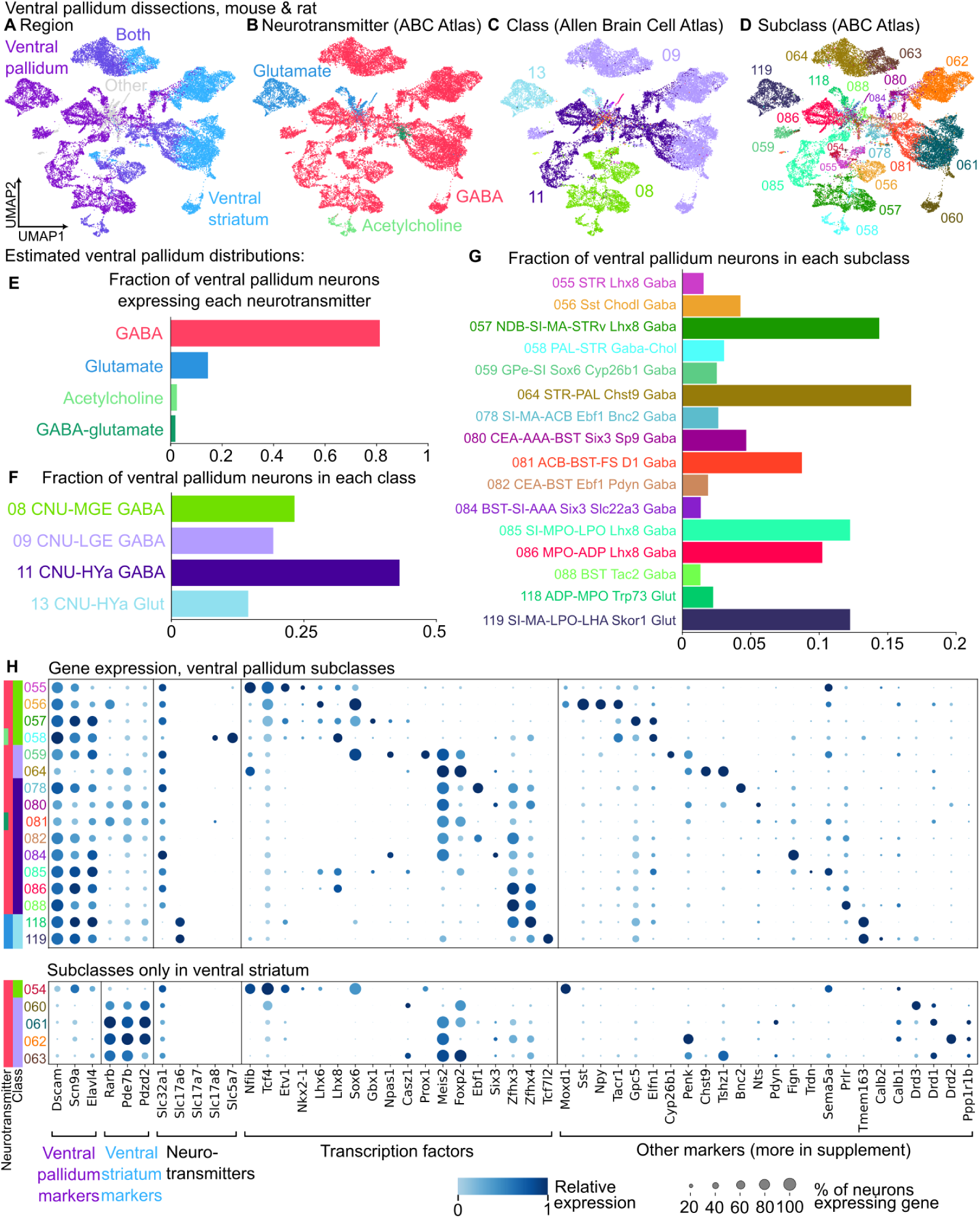
Myriad genetic cell type markers among VP neuronal subclasses. (A) Neurons from VP dissections in mouse and rat, colored by inferred brain region of origin. (B) As in (A), colored by ABC Atlas neurotransmitter assignment. (C) As in (A), colored by ABC Atlas class. (D) As in (A), colored by ABC Atlas subclass. (E) Estimated distribution of neurotransmitter expression in VP neurons. (F) Estimated distribution of ABC Atlas class in VP neurons. (G) Estimated distribution of ABC Atlas subclass in VP neurons. (H) For each neuronal subclass, the fraction of neurons expressing (size) and relative mean expression (color) of marker genes of interest. Plotted for the neurons from VP dissections in mouse and rat.

Our dataset revealed the expression patterns of many cell type markers in VP, both canonical and novel (Fig. 3H). First, by comparing expression levels in all VP and STRv neurons from the VP dissections, we sought markers that best differentiated the two regions. We found that *Dscam*, *Scn9a*, and *Elavl4* were highly expressed in VP but not STRv, and *Rarb*, *Pde7b*, and *Pdzd2* were highly expressed in STRv but not VP. These markers could be useful for anatomically differentiating STRv and VP in subsequent studies.

The expression of neurotransmitter transporter genes highlighted interesting findings. Confirming previous reports, the primary glutamatergic subclasses in VP expressed Vglut2 (*Slc17a6*), and there were no subclasses expressing Vglut1 (*Slc17a7*) (*22*). A subset of cholinergic neurons, in addition to the choline transporter (*Slc5a7*), expressed Vglut3 (*Slc17a8*). The remaining Vglut3 neurons were members of the 081 ACB-BST-FS D1 Gaba subclass, located at the border of VP, STRv, and BNST, and co-expressed GABA (*Slc32a1*) markers, constituting the GABA-glutamate co-releasing neurons mentioned above.

Transcription factor expression further delineated VP neuronal subclasses. All MGE sub-classes, as well as two from the HYa GABA class, expressed *Nkx2.1* and its downstream partners, *Lhx6* and *Lhx8* (also known as *Lhx7*), an established developmental program for pallidal neurons and striatal interneurons (*31–33*). These subclasses included well-known striatal interneuron types, including 054 STR Prox1 Lhx6 Gaba (parvalbumin-expressing, Fig. S5), 056 Sst Chodl Gaba (somatostatin), and 058 PAL-STR Gaba-Chol (cholinergic), as well as three of the major GABAergic subclasses in VP: 057 NDB-SI-MA-STRv Lhx8 Gaba, 085 SI-MPO-LPO Lhx8 Gaba, and 086 MPO-ADP Lhx8 Gaba. Npas1 labeled a population of GABAergic neurons mostly distinct from Nkx2.1, including 059 GPe-SI Sox6 Cyp26b1 and and 084 BST-SI-AAA Six3 Slc22a3 (*33; 34*). Other factors of interest included *Nfib*, *Etv1*, *Gbx1*, *Casz1*, *Prox1*, *Foxp2*, and *Tcf7l2* (*35; 36*), all with unique expression patterns.

The distribution of other markers often used in VP (*22; 35*) varied from highly restricted in distinct clusters to very broadly expressed. Among the restricted markers, somatostatin (*Sst*) and neuropeptide Y (*Npy*) were predominantly expressed in the 056 Sst Chodl Gaba cluster. *Cyp26b1* selectively labeled 059 GPe-SI Sox6 Cyp26b1 Gaba cluster. *Chst9* exclusively labeled the largest subclass in VP, 064 STR-PAL Chst9 Gaba, which is also present in NAc and BNST (Figs. 2K-L). *Bnc2* selectively labeled 078 SI-MA-ACB Ebf1 Bnc2 Gaba neurons. *Tmem163* expression was most prevalent in the two glutamatergic subclasses. These markers could be used to target populations of transcriptionally similar neurons to study in VP.

Other commonly used markers were more broadly expressed across subclasses. *Tacr1* (the receptor for substance P, the anatomical marker for VP), neurotensin (*Nts*), *Sema5a*, calretinin (*Calb2*), calbindin (*Calb1*), and the dopamine receptors (*Drd1*, *Drd2*, and *Drd3*), as well as a variety of genes involved in neurotransmitter, hormone, opioid, and cannabinoid signaling (Fig. S6), were widely distributed. Two markers each used to functionally interrogate VP neuronal function, enkephalin (*Penk*) and parvalbumin (*Pvalb*, see Fig. S5) (*15; 24*), were expressed across many GABAergic subclasses, suggesting they do not correspond to transcriptionally distinct VP subpopulations.

Given the prevalence of VP research in both mice and rats, it was important to validate these expression patterns in both species. Plotting the marker genes for nuclei from mice or rats revealed nearly identical gene expression patterns across neuronal subclasses regardless of species (Fig. S7), indicating strong rodent homology for VP neuron identities.

### High-fat diet alters oxidative phosphorylation and inhibitory signaling genes

With its known involvement in feeding behaviors (*21; 22*), there is interest in how VP may contribute to long-term dietary impacts (*37*). We used a high-fat diet model to explore widespread changes in gene expression accompanying diet-induced weight gain. Groups of 6 mice (3 male, 3 female) were placed on high-fat diet for 2 or 5 weeks, matched diet for 5 weeks, or maintained on standard chow for 2 weeks before tissue collection (Fig. 4A). At each time point, mice on high-fat diet gained more weight compared to control mice (Fig. 4B). We looked for changes in gene expression across all major cell types, dividing neurons into VP GABA classes, VP Glutamate, STRv D1 (subclass 061), and STRv D2 (subclass 062) (Fig. 4C). We detected differentially expressed genes (DEGs) (5 w high-fat compared to 5 w matched diet and 2 w standard chow) in all cell types (Fig. 4D). We trained classifiers to decode high-fat cells from matched diet cells within each cell type, all of which outperformed shuffled data with comparable accuracy (Fig. 4E), suggesting that the magnitude of the diet impact was similar for all cell types. The change in expression from 2 w standard to 2 w high-fat or 5 w high-fat was highly correlated (Fig. 4F) such that genes upregulated at 5 w were also typically upregulated at 2 w, with same true for downregulated genes (Fig. 4G), indicating a consistent impact of diet across time points.

**Figure 4.**
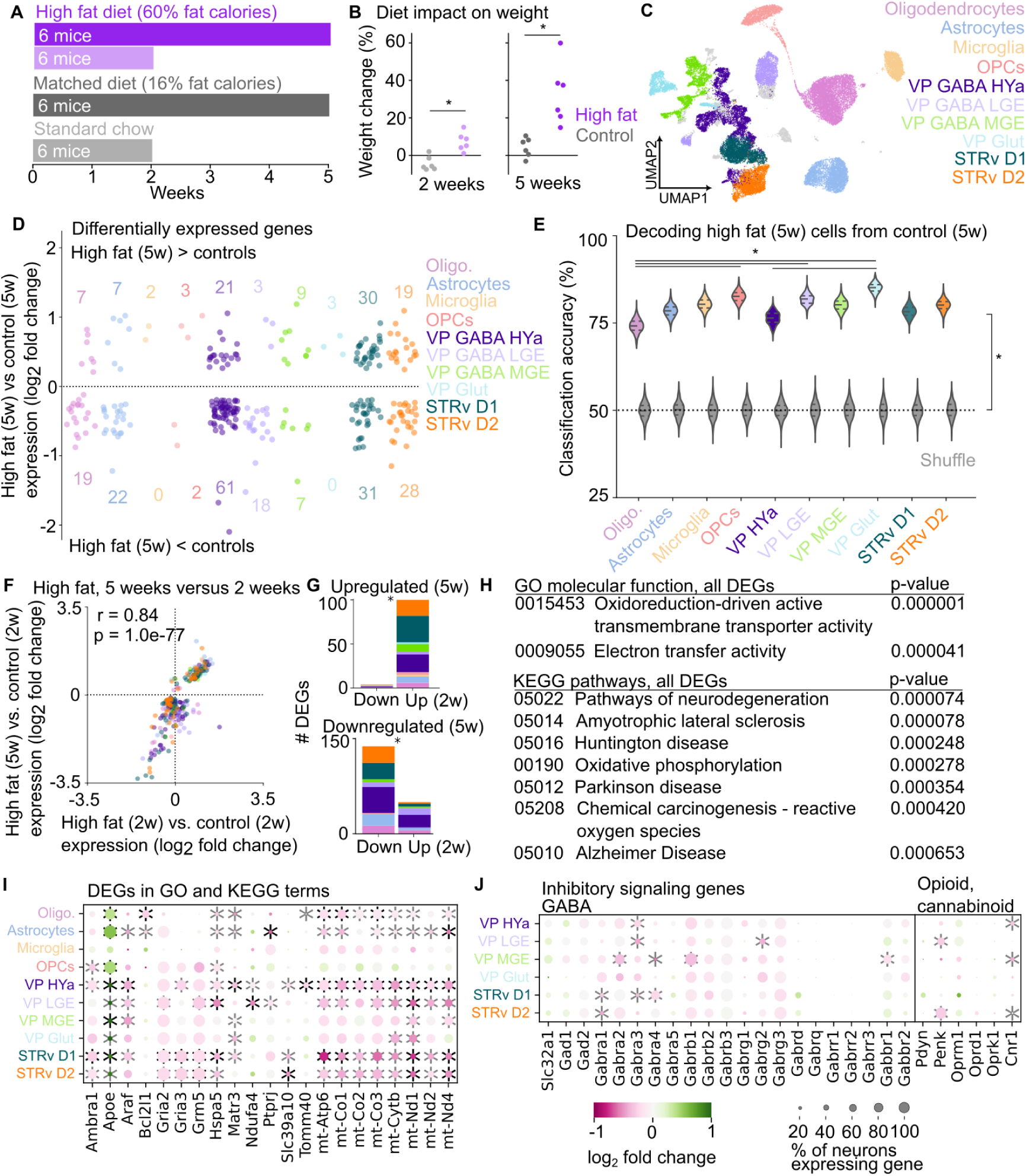
High fat diet alters oxidative phosphorylation and inhibitory signaling genes. (A) Diet manipulation for mice included in the VP tissue dissections. (B) Change in weight for each mouse following diet exposure. 2 w: *t* = *−*4.59, *p* = 0.001, 5 w: *t* = *−*4.06, *p* = 0.002. (C) Cell types used for high-fat diet analysis. (D) The log_2_ fold change in expression, 5 w high-fat compared to matched diet, for all differentially expressed genes (DEGs) from each group. (E) Classifier performance decoding high-fat from matched diet cells from each cell type using expression of all DEGs from (D), bootstrapped, 330 cells per group, 1000 repetitions, with replacement. Also with diet labels shuffled. * denotes *p <* 0.05, Bonferroni-corrected. (F) The log_2_ fold change in expression, 5 w high-fat compared to standard chow versus 2 w high-fat compared to standard chow, for all DEGs, with Pearson’s r and p-value denoted. (G) For all DEGs increased (top) or decreased (bottom) at 5 w compared to control diets, the number with log_2_ fold change below (‘Down’) or above (‘Up’) 0 at 2 w compared to standard chow. Upregulated: *χ*^2^ = 88.6, *p* =4.8e-21, downregulated: *χ*^2^ = 41.2, *p* =1.3e-10. (H) GO and KEGG terms with significant overlap with the list of DEGs from (D). (I) The log_2_ fold change in expression, 5 w high-fat compared to matched diet, for DEGs present in the GO and KEGG terms in (H), plotted for all cell types and sized by % neurons expressing the gene in that cell type. * denotes DEG (see Methods). (J) The log_2_ fold change in expression, 5 w high-fat compared to matched diet, for genes involved in GABA, opioid, and cannabinoid signaling with substantial expression in our dataset (Fig. S6), plotted for all neuronal cell types and sized by % neurons expressing the gene in that cell type. * denotes DEG (see Methods).

To perform an unbiased analysis of gene pathways altered by high-fat diet, we input all DEGs into the GO and KEGG databases (*38; 39*). This approach revealed a significant over-representation of genes related to oxidative phosphorylation, which also plays a role in many neurodegenerative diseases (Fig. 4H-I). Due to evidence of altered inhibitory signaling in VP following high-fat diet (*26; 37*), as well as a specific role of VP opioid signaling in feeding (*21; 27*), we looked more closely at genes related to GABA, opioid, and cannabinoid signaling (Fig. 4J). We found that a number of GABA receptor subunits were downregulated in VP GABAergic neurons, as well as cannabinoid receptor 1 (*Cnr1*). Although we detected no significant changes in opioid receptor expression, the gene for enkephalin (*Penk*) was downregulated in the neuronal populations with the largest *Penk* expression: VP GABAergic neurons from lateral ganglionic eminence and STRv D2 neurons, which release enkephalin in VP (*22*). These results encourage continued examination of the relationship between high-fat diet and alterations to inhibitory signaling mechanisms in this circuit.

## Discussion

Our experiments revealed the molecular diversity of VP cell types. By sequencing individual nuclei from mouse and rat VP tissue and integrating our results with a standardized atlas, we thoroughly characterized rodent VP neuron subtypes and contextualized them within brain-wide expression patterns. The large number of GABAergic subclasses (14) and their spatial distribution beyond traditional VP borders will be important considerations for future cell type-specific experiments in VP and adjacent brain regions. Our results highlight the lack of region-specific transcriptionally-defined cell types in VP and beyond, as well as the incomplete overlap between classic cell type markers and transcriptionally distinct neuron groups.

Leveraging the ABC Atlas (*25*) and the MapMyCells tool (which we validated with our own clustering integration method, Fig. S3), we were able to assign atlas labels to neurons in our (and other) datasets. This had many advantages over more traditional classification approaches. First, it ensured a consistent level of granularity when dividing neurons into subtypes. For instance, some cell types were too rare in VP to reliably cluster separately when only considering VP neurons, but they were more clearly distinguished in the context of additional neurons from other brain regions (for example, in Figs. 2I and S3E). Second, it allowed us to look at the spatial distribution of neurons with similar expression patterns (i.e. members of the same subclass) in the integrated MERFISH datasets. Third, it will provide consistent terminology for the same cell types across future studies.

By specifically dissecting the overlap between VP cell types and neighboring regions, we took a crucial step towards an improved understanding of VP regional identity. With sequencing data alone, it is unclear whether neurons expressing markers that label neurons from neighboring regions is indicative of overlapping cell types between regions or ‘contamination’ from nearby tissue during dissection. By combining spatial MERFISH data and existing RNA-sequencing datasets from neighboring regions, we validated the wide spatial distribution of the subclasses we identified in VP, extending into neighboring striatal, extended amygdalar, and hypothalamic regions. This finding highlights the importance of considering cell types in each region within the context of the whole brain; it may be incorrect to assume that a transcriptionally-defined cell type from a given region is unique to that region. Also, by demonstrating that VP shares cell types with regions well outside the basal ganglia, our experiment provides additional evidence that VP is not simply a basal ganglia nucleus but perhaps a transition area between multiple regions. Our study raises important questions about the interaction between genetic underpinnings and anatomical connectivity in defining a region.

Our sequencing results were consistent with previous reports of VP neuronal diversity, starting with the presence of neurons releasing GABA, glutamate, or acetycholine. Of the three neurotransmitter classes, cholinergic neurons are known to make up the smallest fraction of VP neurons (*22; 23*), with one microscopy study estimating the proportion to be 11% (*17*). In our sequencing results, only 2.4% of VP neurons were cholinergic. Perhaps the larger size of cholinergic neuron cell bodies (*22*) explains the discrepancy between these methods. Our data confirmed previous reports about the presence of cholinergic neurons expressing Vglut3; interestingly, this population selectively innervates the basolateral amygdala (*40*). Additionally, we confirmed high levels of expression of *Tacr1*, the receptor for substance P, in cholinergic neurons (*41*).

The largest subclass (GABAergic or otherwise) in VP was 064 STR-PAL Chst9 Gaba, which was also present in STRv and was very selectively labeled by *Chst9*. This subclass, along with the STRv-only subclass 063 STR D1 Sema5a, was also labeled by *Tshz1*, which has been useful for identifying a functionally distinct population of striatal neurons (*42*). The presence of *Chst9* neurons in both VP and STRv makes it a particularly interesting population to study in the future because it contrasts with the major STRv GABAergic subclasses that are selectively localized to striatum (Fig. 2E). We were able to confirm that this subclass has the highest expression of *µ*-opioid receptor (Fig. S3B), echoing a report from NAc (*43*). Another subclass of interest that spanned both regions (as well as BNST) was 081 ACB-BST-FS D1 Gaba, which also contained the dual GABA-Glutamate neurons, which could warrant additional investigation.

The expression of transcription factors across the subclasses we identified from our VP dissections helped link these subclasses to established literature on pallidal development (Fig. 3H) (*31–33*). *Nkx2.1* was expressed in all MGE subclasses detected in our VP dissections, including the striatal interneuron subclasses, as well as 085 SI-MPO-LPO Lhx8 Gaba and 086 MPO-ADP Lhx8 Gaba, which originate in preoptic area. There were some differences in the expression of the downstream partners of *Nkx2.1*, *Lhx6*, and *Lhx8*, with *Lhx6* more prominent in 056 Sst Chodl Gaba (somatostatin-expressing neurons) and the striatal-only subclass (putatively parvalbumin interneurons, Fig. S5), and *Lhx8* more prominent in 058 PAL-STR Gaba-Chol (cholinergic neurons) and 086 MPO-ADP Lhx8 Gaba. *Sox6* expression was generally consistent with the finding that it acts downstream of *Lhx6* (*44*). One interesting implication of these results is that many STRv interneurons, including those expressing somatostatin and acetylcholine, are transcriptionally and developmentally similar to neurons in VP, sharing the same subclass and transcription factor expression and detected in dissections from both regions (Fig. S4C).

As reported for globus pallidus (*33; 34*), the transcription factor *Npas1* labeled a population of GABAergic neurons mostly distinct from *Nkx2.1* and *Lhx6*, which in our experiment included 059 GPe-SI Sox6 Cyp26b1 and 084 BST-SI-AAA Six3 Slc22a3 Gaba. One notable feature of the *Npas1* population is its axonal projections back to striatum (*45; 46*). Another factor, *Gbx1* labeled two of the largest GABAergic subclasses in VP, 057 NDB-SI-MA-STRv Lhx8 Gaba and 085 SI-MPO-LPO Lhx8 Gaba, making it a potentially useful target for functional manipulations. *Tcf7l2* selectively labeled the largest subclass of glutamatergic neurons, 119 SI-MA-LPO-LHA Skor1 Glut, consistent with a report of its importance in maintaining glutamatergic identity in thalamus (*47*). Expression of *Zfhx3* and *Zfhx4* was high across all anterior hypothalamic subclasses, both GABAergic and glutamatergic.

Interestingly, many of the cell type markers commonly used in VP do not correspond to transcriptionally distinct populations of neurons. Enkephalin (*Penk*) has been shown to label a functionally distinct population of GABAergic neurons in VP (*24*), but it was expressed across many GABAergic subclasses in our experiment (Fig. 3H). Another frequently used marker is parvalbumin (*15; 16*). Although undetected in our dataset, it was present in the PAL dissections from the ABC Atlas, possibly reflecting a difference between nucleus and whole cell sequencing, so we analyzed its expression pattern among the same subclasses in that dataset. Parvalbumin was most highly expressed in 057 NDB-SI-MA-STRv Lhx8 Gaba and 059 GPe-SI Sox6 Cyp26b1 Gaba, but it was rather widely expressed overall, particularly in contrast to STRv, where it was restricted to 054 STR Prox1 Lhx6 Gaba (Fig. S5A). Therefore, although it has been useful as a marker in VP like enkephalin, parvalbumin expression does not correspond to a transcriptionally uniform VP subpopulation. Subsequent work could interrogate separate functional roles for parvalbumin neurons belonging to distinct subclasses.

Building upon evidence that VP is altered by exposure to high-fat diet (*26; 27*), controls preference for fatty foods (*48*), and modulates weight gain from high-fat diet (*49*), we investigated how prolonged high-fat diet changes widespread gene expression in VP. We detected DEGs in all cell types when comparing gene expression after 5 w of high-fat diet to 5 w of matched diet and to 2 w of standard diet. The statistical power to detect DEGs depends on the number of cells, which varied across groups, so we opted not to interpret the total number of DEGs in each group. Instead, we used a matched number of cells from each group to decode diet condition, revealing a similarly detectable impact of diet in all cell types (Fig. 4E). In fact, VP Glutamate neurons were classified with higher accuracy than VP GABA HYa neurons despite having fewer DEGs. Future experiments could increase the number of cells to powerfully and rigorously identify additional DEGs in the VP Glutamate population. Interestingly, the 119 SI-MA-LPO-LHA Skor1 Glut subclass had the highest expression of the GLP-1 receptor (Fig. S6B), a known regulator of feeding behavior (*50*), providing additional incentive to study a role for VP Glutamate neurons in overeating.

The majority of the DEGs that were members of the significant GO and KEGG terms were related to metabolic processes, particularly mitochondrial DNA-encoded genes involved in the function and maintenance of the electron transport chain (Fig. 4I). Notably, we observed a robust increase in expression of the *Apoe* chylomicron protein across almost all clusters. Several genes involved in regulation of apoptosis (*Araf*, *Bcl2l1*, *Hspa5*) were also altered. The overall pattern of gene expression indicates that high fat diet induced a metabolic shift within many of the cell type clusters we observed, with oligodendrocytes, astrocytes, two VP GABA sub-classes, and striatal D1 and D2 neurons displaying similar trends. Many of these DEGs are also implicated in various neurodegenerative disease pathways (Fig. 4H). Further investigation will be needed to determine whether high-fat diet induced metabolic changes in this circuit play a role in neurodegeneration.

We also took a more targeted approach to look at mechanisms previously described in the literature through which high-fat diet can impact VP function. Exposure to high-fat diet lowers firing rate in VP and, interestingly, mice with the propensity to gain the most weight have potentiated inhibitory signaling onto VP neurons (*26*). This encouraged us to look more closely at genes related to inhibitory mechanisms. We found downregulation of a number of GABA receptor subunits in VP GABA classes (Fig. 4J), which is difficult to interpret in the context of sequencing data alone, but it could indicate adaptations to altered GABAergic input. Another prior finding was increased *δ*-opioid receptors in VP following prolonged high-fat diet exposure (*27*), a possible mechanism for potentiated inhibitory input (*37*). We found low expression of the *Oprd1* gene overall (Fig. S6), hindering our ability to detect a similar change in our experiment. We did, however, see reductions in enkephalin expression in both VP and NAc neurons, suggesting that opioid signaling may indeed be altered in this circuit following high-fat diet exposure. We also saw reduced cannabinoid receptor (*Cnr1*) expression in VP, another inhibitory input and a known regulator of feeding (*51*) implicated in transgenerational dietary impacts (*52*), suggesting that VP GABA neurons may be an important site to study cannabinoids’, and opioid-cannabinoid interactions’ (*53*), involvement in long-term feeding adaptations.

Overall, our work builds upon research demonstrating a role for VP that extends beyond its original characterization as a basal ganglia nucleus. We have shown here that VP contains a collection of cell types from brain regions not typically associated with the basal ganglia, suggesting that VP is more of a transitional area, combining striatal input with the functional capabilities of adjacent regions to robustly influence motivated behavior. Future work can build upon our results by examining the interplay between molecular identity and anatomical connectivity in defining the role each cell type plays in reward, feeding, and other contexts.

## Acknowledgments

Thank you to Nailyam Nasirova for assisting with the rats. This work was supported by National Institutes of Health grants F32DA053714 (D.J.O.), R37DA032750 (G.D.S.), R01DA054317 (G.D.S and S.M.F.), and P30DA048736 (G.D.S.).

## Author contributions

Conceptualization: D.J.O., R.C.S., G.D.S.; data collection: D.J.O., R.C.S., A.J.B., G.D.S.; data curation: D.J.O., R.C.S., C.T.B.; data interpretation: D.J.O., R.C.S., C.T.B., A.J.B., G.D.S.; formal analysis: D.J.O.; visualization: D.J.O.; writing - original draft: D.J.O., C.T.B.; writing - review & editing: D.J.O., R.C.S., C.T.B., A.J.B., S.M.F., G.D.S.; funding acquisition: S.M.F., G.D.S..

## Data and code availability

The data and code for this manuscript will be publicly available by the time of publication.

## Methods

### Subjects

Subjects included mice (*n* = 24) and rats (*n* = 4). Mice were male (*n* = 12) and female (*n* = 12) C57BL/6 strain group-housed on a reverse 12hr light/dark cycle. Rats were male (*n* = 2) and female (*n* = 2) Sprague-Dawley strain pair-housed on a reverse 12hr light/dark cycle. All experimental procedures were performed in strict accordance with protocols approved by the Animal Care and Use Committee at the University of Washington.

### Diet manipulation

We had four experimental groups of mice. Each group contained two cages, with 3 males or 3 females in each. Two groups had ad lib access to high-fat diet (60% calories from fat, Bio-Serv F3282) for either 2 or 5 weeks. One group had ad lib access to a matched control diet (16% calories from fat, Bio-Serv F4031) for 5 w. The final group was maintained on standard chow for 2 w. Mice were age-matched such that they were P76 - P90 at time of tissue collection, with a mean of P83 - P84 for all groups.

### Tissue collection

For experiments utilizing mice, all mice were anesthetized with isoflurane and transcardially perfused with ice-cold NMDG buffer. Brains were extracted and sliced on a fresh-tissue vibratome. Typically, 4 tissue slices of 300 *µ*M thickness were collected for bilateral VP microdissection with a scalpel, resulting in 8 VP tissue samples. Samples were flash-frozen on dry ice and transferred to -80 C for storage. For experiments utilizing rats, rats were injected with 0.7 mL Euthasol for terminal anesthesia, and then perfused with ice-cold NMDG buffer. VP tissue samples were collected using the same procedure employed for mice excepting for with 400 *µ*M slices. We also collected 8 NAc tissue samples from the same rats. Example dissections are depicted in Figs. 1A and S1.

### Nuclear isolation

VP tissue samples from each experimental group (mice) and from all rats were pooled together for a total of 5 nuclear isolation batches by combining samples in a glass Dounce homogenizer containing lysis buffer. We did the same for NAc tissue samples from rats for an additional nuclear isolation batch. After homogenization and a five-minute incubation, samples were diluted using wash buffer, passed through a 40 uM cell strainer, and centrifuged at 500 x g for 10 minutes at 4 C to remove debris and pellet nuclei. This pellet was resuspended in a homogenate solution and combined 1:1 with a 50% iodixanol solution. To further clean up the nuclei pellet using density gradient centrifugation, we underlaid the resulting 25% iodixanol layer with a 30% layer and centrifuged at 3000 x g for 15 minutes at 4 C. Finally, the nuclei pellet was resuspended in wash buffer and nuclei were assessed for yield and quality with a hemocytometer (INCYTO, #DHC-N01) and a fluorescent microscope (Zeiss ApoTome.2). For additional detail see (*54*).

### Library preparation and sequencing

We used the 10X Genomics Chromium 3’ NextGEM kit v3.1 (Dual Index) to generate cDNA libraries from each nuclear isolation batch. Isolated nuclei preparations were diluted to a concentration of 1,000/uL and loaded onto the 10X chip, targeting capture of 10,000 single barcoded nuclei. We loaded rat nuclei preparations into two lanes on the chip, resulting in two VP and two NAc reactions. We then prepared libraries for sequencing. Sequencing was performed on the NovaSeq 6000 platform to a depth of 350M paired reads/reaction (mice) or the NovaSeq X Plus platform to a depth of 750M paired reads/reaction (rats) by Novogene Co. We aligned the sequencing results to reference genomes using the 10x Genomics Cloud Analysis tool, using their provided mouse reference genome, Mouse (mm10) 2020-A, and creating our own rat reference genome with 10x Genomics Cell Ranger software. FASTA and GTF files for Rattus norvegicus (mRatBN7.2) were downloaded from Ensembl.

### Single cell analysis

The resulting cells x genes matrices were loaded into Python for further analysis, using packages Scanpy (RRID:SCR 018139) (*55*) and scvi-tools (*56*). First, we removed cells with fewer than 1,000 and greater than 20,000 (mice) or 40,000 (rat) unique molecular identifiers (UMIs), as well as cells with greater than 10% or 4% of UMIs from mitochondrial or ribosomal genes, respectively. We then removed any genes that were present in no remaining cells. We fit an scVI model (*57*) with default settings on the 2,000 most variable genes (Scanpy ‘highly variable genes’ with ‘seurat v3’ and batch correction options) to find expression patterns across all mouse reactions and all rat reactions, correcting for batch. We then fit Solo (*58*) models on each batch to detect and remove doublets. We then used scVI model latent variable values for the remaining cells to calculate nearest neighbors (Scanpy ‘neighbors’ with n neighbors = 50) and generated a UMAP (Scanpy UMAP with min dist = 0.5) using these connectivities. We divided cells into clusters according to these connectivities (Leiden algorithm) and assigned them to cell types guided by the expression of canonical markers (Fig. S2). We followed the same procedure to identify neurons in datasets from NAc (*28*), BNST (*29*), POA (*30*), and PAL (*25*).

To combine neurons from multiple datasets (neurons from VP mouse and rat: Fig. 1C, neurons from VP, NAc, BNST, and POA: Fig. S4A, neurons from VP, NAc, BNST, and POA in VP subclasses: Fig. 2I, neurons from VP, NAC, and PAL: Fig. S3B) we used scVI with recommended settings for integrating datasets (n layers = 2, n latent = 30, gene likelihood=‘nb’), correcting for batch, UMIs per cell, experiment, and species. The latent variables from these scVI models were used to generate nearest neighbors (n neighbors = 15) for UMAP (and clustering in Fig. S3).

### Atlas integration

We used the Allen Brain Cell (ABC) Atlas (*25*) tool MapMyCells (RRID:SCR 024672) to assign class, subclass, supertype (and associated neurotransmitter), and cluster to neurons from each dataset. We validated this tool for neurons from VP and NAc by clustering neurons from VP, NAc, and PAL (using scVI integration detailed above) with Leiden algorithm (resolution = 0.2) and then, for each cluster, clustering again (resolution = 1.5 for 10 largest clusters, 1 for remaining) and assigning neurons from VP and NAc dissections to the majority label from the PAL neurons within that subcluster (PAL labels were assigned in the generation of the ABC Atlas). The two approaches yielded nearly identical results (Fig. S2H).

### Validation with MERFISH

The ABC Atlas contains several MERFISH experiments that detail spatial information about cells that have been assigned Atlas labels (*25*). To ascertain the spatial distribution of sub-classes we detected in our sequencing data, we analyzed 4 experiments containing VP sections in the Mouse Whole-Brain Transcriptomic Cell Type Atlas and the MERFISH Whole Mouse Brain datasets (https://portal.brain-map.org/atlases-and-data/bkp/abc-atlas). First, we visually inspected the spatial distribution of all subclasses we identified in our VP dissections to verify their location in and around VP. To find the number of VP neurons in each subclass (Fig. 2B), we selected the substantia innominata and magnocellular nucleus spatial filters, counted the number of cells belonging to each subclass of interest, and divided by the total number of cells belonging to neuronal subclasses. We used the Zhuang-ABCA-1 experiment to produce example images of region identity and subclass distribution in Fig. 2.

### Estimation of VP proportions

The presence of STRv-only subclasses in our VP dissections indicated a substantial proportion of STRv neurons we needed to account for when estimating the number of VP neurons belonging to subclasses present in both VP and STRv. We used the ratio between neurons in STRv-only subclasses (054, 060, 061, 062, 063) in NAC experiments and neurons in those same subclasses in our VP dissections to estimate the fraction of neurons in the VP/STRv subclasses (055, 056, 058, 064, 078, 081) that were actually from VP in our VP dissections. Using these subclass estimates, we populated class and neurotransmitter estimates accordingly.

### Gene expression analysis

For subsequent gene expression analysis, we first normalized the number of UMIs per nucleus to 10,000. To help identify marker genes in VP, we used the Scanpy function ‘rank genes groups’ (with method = ‘wilcoxon’) on log-converted counts (‘log1p’ function) for all neurons in VP subclasses. To find maximally differentially expressed genes (DEGs) between VP and STRv, we used ‘rank genes groups’ (with method = ‘wilcoxon’) on log-converted counts (‘log1p’ function) comparing neurons in VP (but not STRv) subclasses to neurons in STRv (but not VP) subclasses. To visualize marker gene expression across subclasses (Figs. 3H, S5, S6, S7), we used the ‘dotplot’ function with standard scale = ‘var’, which normalizes expression from 0 to 1 across the means of all groups (subclasses in this case) for each gene.

### High-fat diet expression analysis

To analyze changes in gene expression induced by high-fat diet, we divided cells into cell types that provided a useful level of granularity while still maintaining sufficient cells per group to detect DEGs. We divided neurons into VP GABA neurons belonging to 08 CNU-MGE GABA, 09 CNU-LGE GABA, or 11 CNU-HYa GABA, VP Glutamate neurons (belonging to 13 CNU-Hya Glut), and STRv neurons belonging to 061 STR D1 Gaba or 062 STR D2 Gaba. To calculate DEGs for each cell type, we first found the log_2_ fold change in mean (normalized) UMIs per condition and then performed a Mann-Whitney U Test to detect a change in expression in the appropriate direction. We then performed a Bonferroni correction, multiplying the p-values by the the number of genes we tested (27,809). To be considered a DEG, there needed to be a significant (*p <* 0.05, corrected) difference between standard chow (2 w) and high-fat diet (5 w) and between matched diet (5 w) and high-fat diet (5 w). To plot the log_2_ fold change, we used the comparison between matched diet (5 w) and high-fat diet (5 w). To compare changes in expression at 2 w and 5 w of high-fat diet, we used the log_2_ fold change at each time point relative to standard chow (2 w). When looking for differences in expression for our targeted gene lists (Figs. 4I-J, we used a Bonferroni correction reflecting the smaller number of genes on those lists. For those figures only, significant DEGs with the stricter cutoff are marked with a black asterisk, and significant DEGs with the looser cutoff are marked with a gray asterisk. To find gene pathways implicated by our broadly detected DEGs, we inputted all DEGs (*n* = 197) into the Scanpy ‘enrich’ function, searching the GO molecular function and KEGG databases. To validate our results, we took 1000 random selections of 197 genes and found the 99th percentile of lowest p-value (*p* = 0.0023 GO, *p* = 0.0019 KEGG) and used this as our adjusted cutoff. Therefore, we estimate a *<* 1% probability that the terms listed are false positives.

We also calculated the ability for a decoder to correctly classify cells from each cell type as belonging to high-fat (5 w) or matched (5 w) diet conditions. For each cell type, we selected 330 cells from each condition (corresponding to number of cells in the condition with the fewest cells across all cell types) with replacement and fit a random forest model (scikit-learn ‘RandomForestClassifier’, n estimators = 30, bootstrap = True) (*59*) to classify cells in each condition, predicting the condition of held-out cells with 5-fold cross-validation. We also did this with condition labels shuffled. We repeated this 1000 times to bootstrap a p-value, using Bonferroni correction for the number of comparisons we made (9 group comparisons with shuffle, 45 comparisons between groups).

### Statistics

All statistical tests were performed with SciPy (*60*) with specific tests and corrections noted in the text. We used Spearman correlation to compare the rankings of most prevalent subclasses across datasets. We used Pearson correlation to compare the degree of gene expression change at 2 w and 5 w of high-fat diet.

---

**Figure S1.**
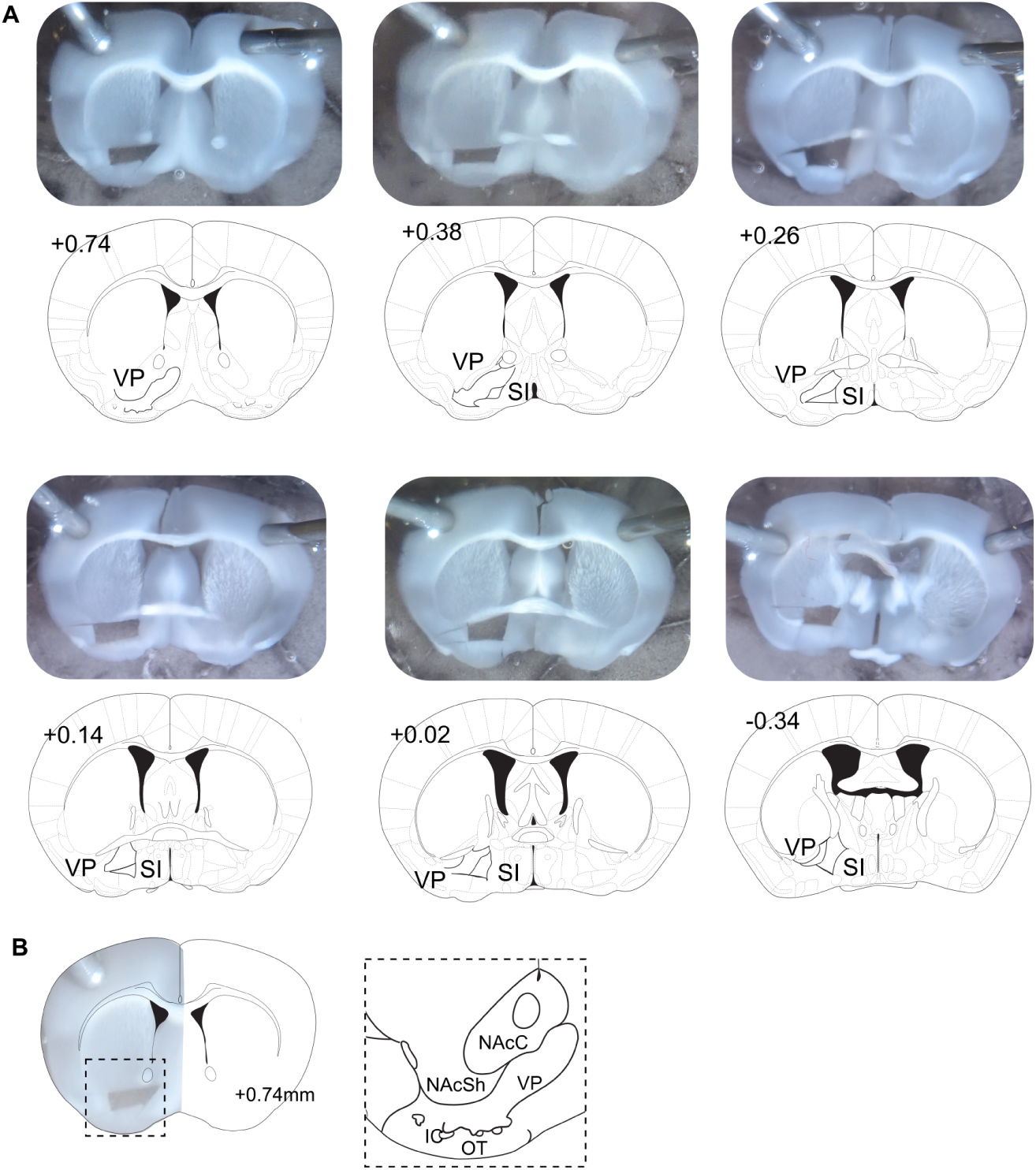
Representative tissue microdissection. (A) Representative VP microdissections from mouse and corresponding Atlas information. (B) Additional detail on a VP dissection with neighboring striatal subregions labeled.

**Figure S2.**
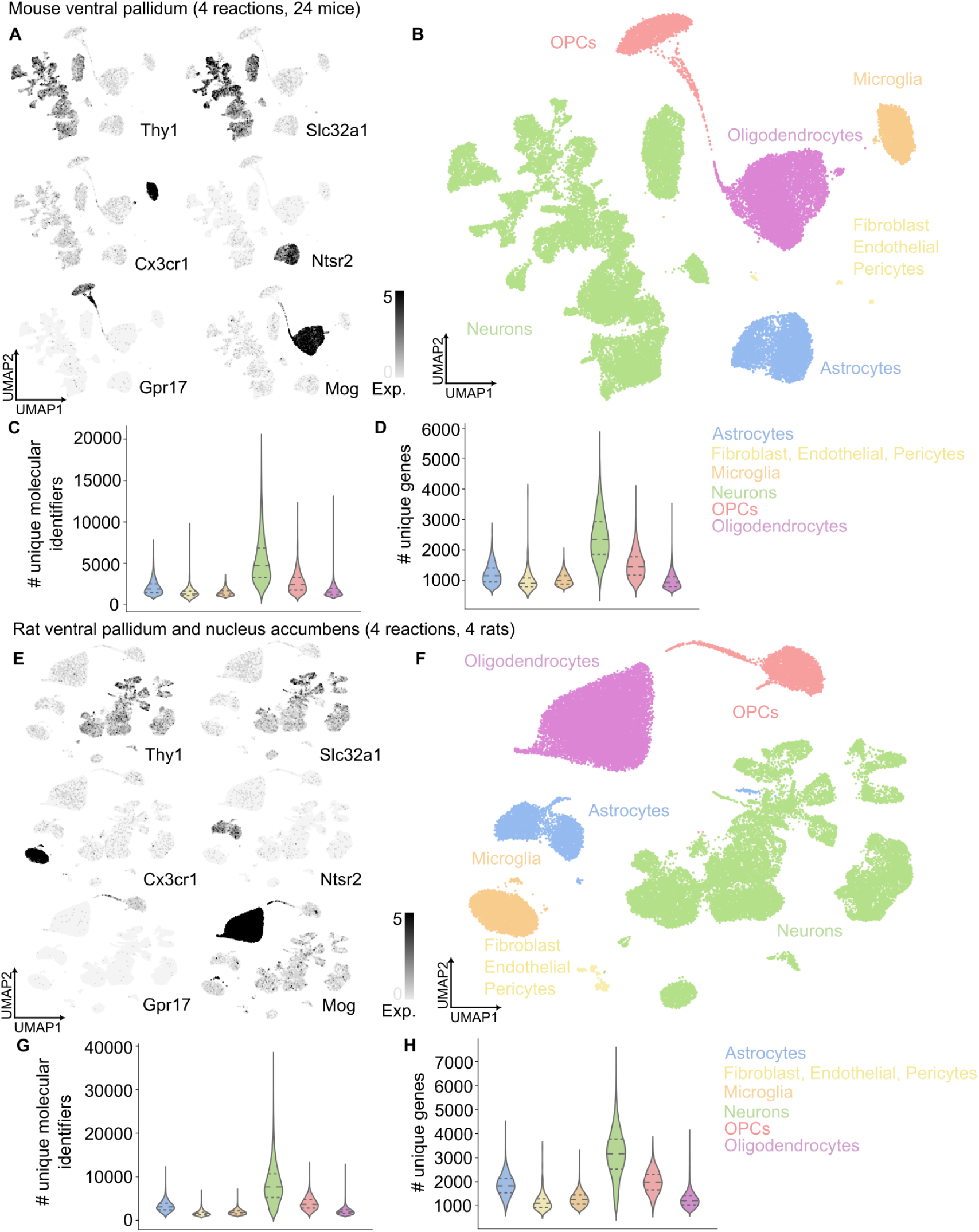
Identification of cell types. (A) Expression of cell type markers among nuclei from mouse VP dissections. (B) Mouse VP nuclei labeled by cell type. (C) For each cell type, the distribution of the number of unique molecular identifiers detected per nucleus. (D) For each cell type, the distribution of the number of unique genes detected per nucleus. (E) Expression of cell type markers among nuclei from rat VP and NAc dissections. (F) Rat VP and NAc nuclei labeled by cell type. (G) For each cell type, the distribution of the number of unique molecular identifiers detected per nucleus. (H) For each cell type, the distribution of the number of unique genes detected per nucleus.

**Figure S3.**
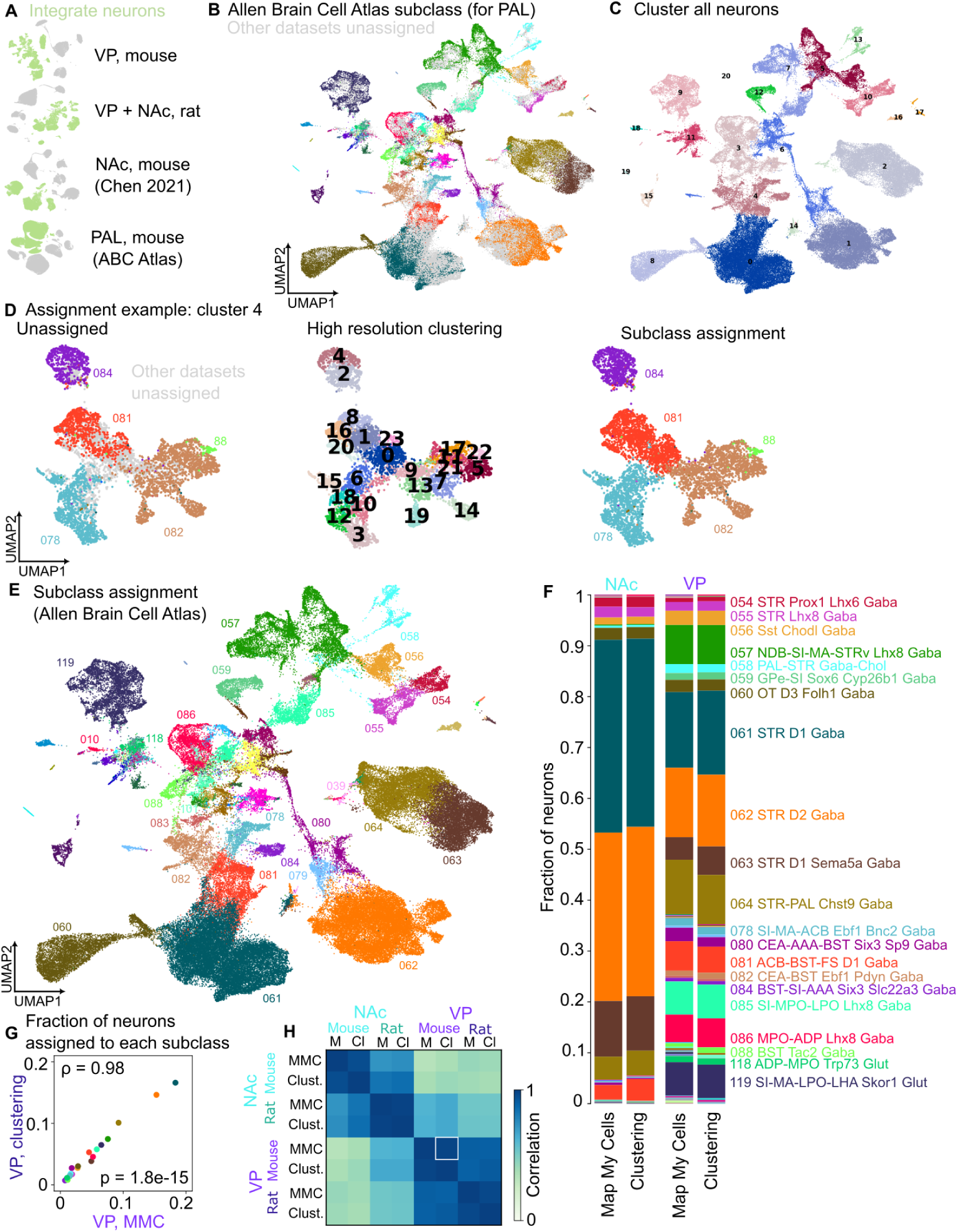
Assignment of Allen Brain Cell Atlas subclasses. (A) UMAP plots of cells from the 4 datasets used for manual Atlas integration, with neurons highlighted in green. (B) Neurons from all 4 datasets, colored by ABC Atlas subclass for the neurons from the ABC Atlas pallidum (PAL) dissections. (C) All neurons divided into clusters with Leiden algorithm. (D) For an example cluster, initial subclass labels (for PAL), additional clustering step, and assignment of subclass labels to remaining neurons in subclass. (E) Subclass assignment for all neurons using the clustering method in (D). (F) Comparison of proportions of NAc and VP neurons assigned to each subclass using MapMyCells (as in main figure) or clustering assignment (as above). (G) The fraction of mouse VP neurons belonging to each subclass, separated for MapMyCells and clustering methods, with Spearman’s *ρ* and p-value denoted. (H) Spearman correlation comparing the fraction of neurons in each subclass for all combinations of dissection region, species, and Atlas integration method. The comparison in (G) is marked with a white box.

**Figure S4.**
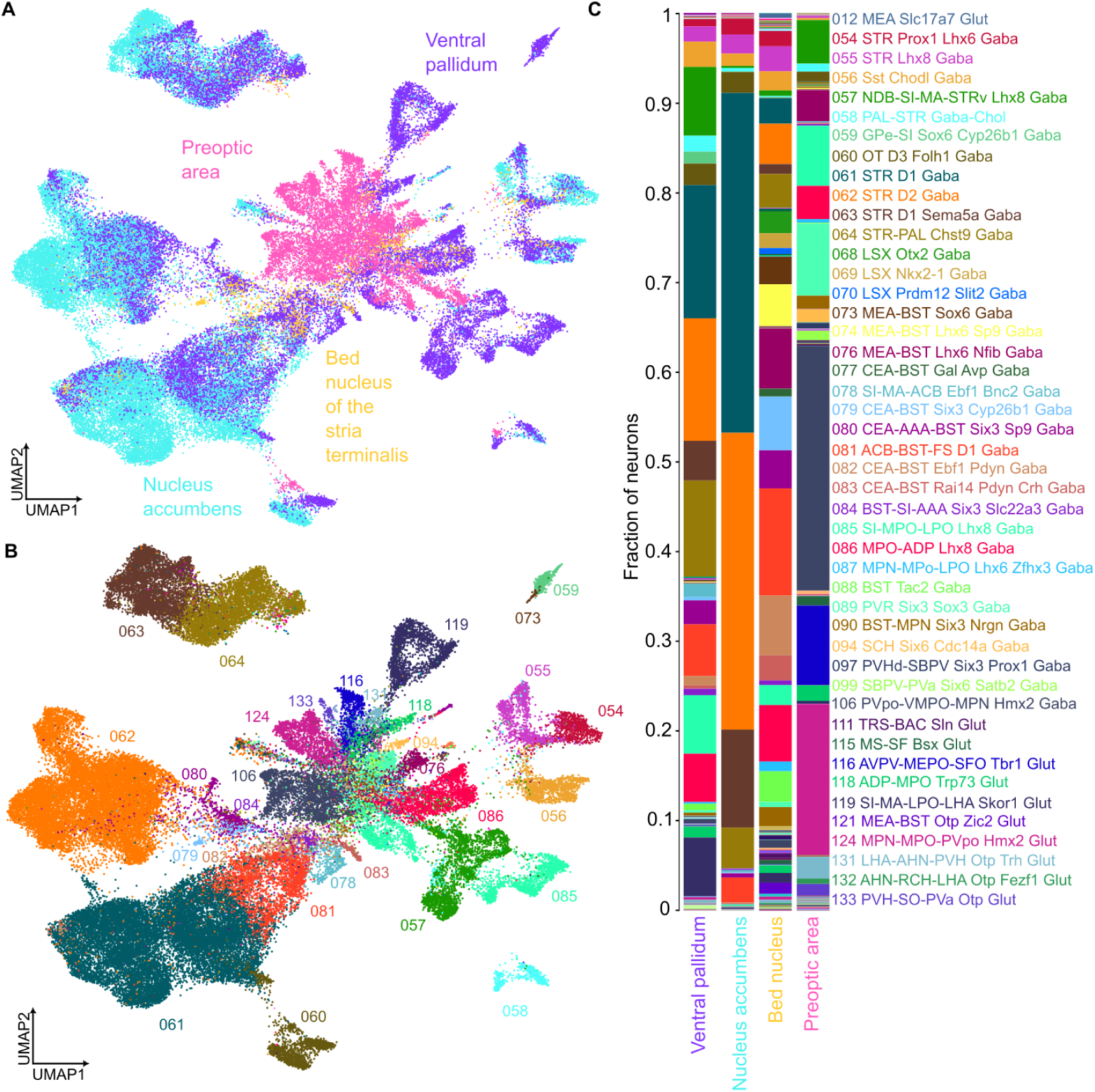
Detail on neurons from four dissection regions. (A) All neurons we analyzed from ventral pallidum, nucleus accumbens, bed nucleus of the stria terminalis, and preoptic area experiments, plotted according to the similarity of their gene expression patterns and colored by dissection region. (B) As in (A), colored by ABC Atlas subclass, as determined by MapMyCells. (C) The fraction of neurons from each dissection region belonging to each subclass. Sub-classes containing *>* 0.5% of neurons from any region are labeled.

**Figure S5.**
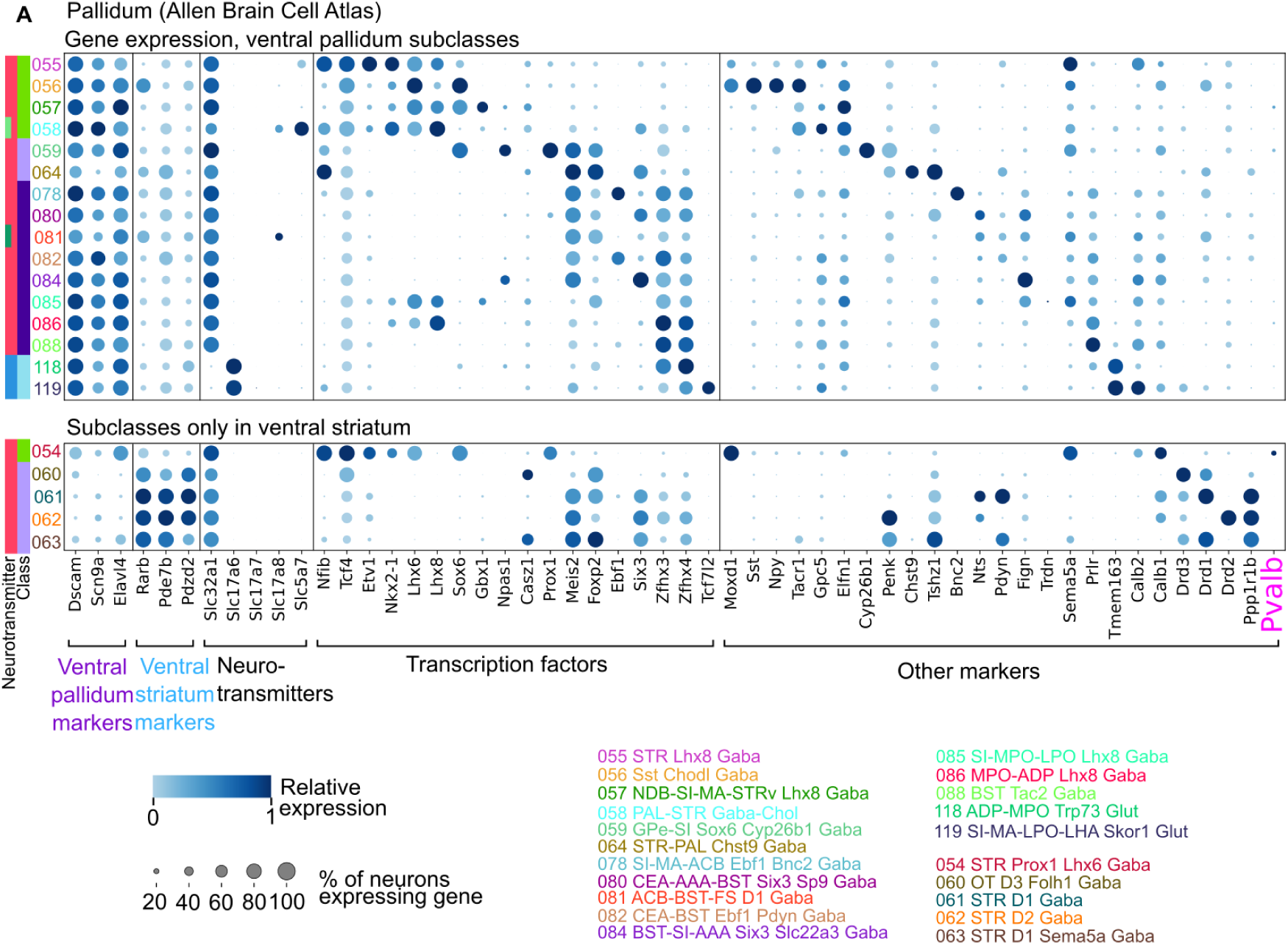
VP expression patterns in Allen Brain Cell Atlas pallidal tissue. (A) For each neuronal subclass, the fraction of neurons expressing (size) and normalized mean expression (color) of marker genes of interest. Plotted for the pallidum neurons from the Allen Brain Cell Atlas. Of note, parvalbumin (Pvalb) is added here because it was better detected in this dataset.

**Figure S6.**
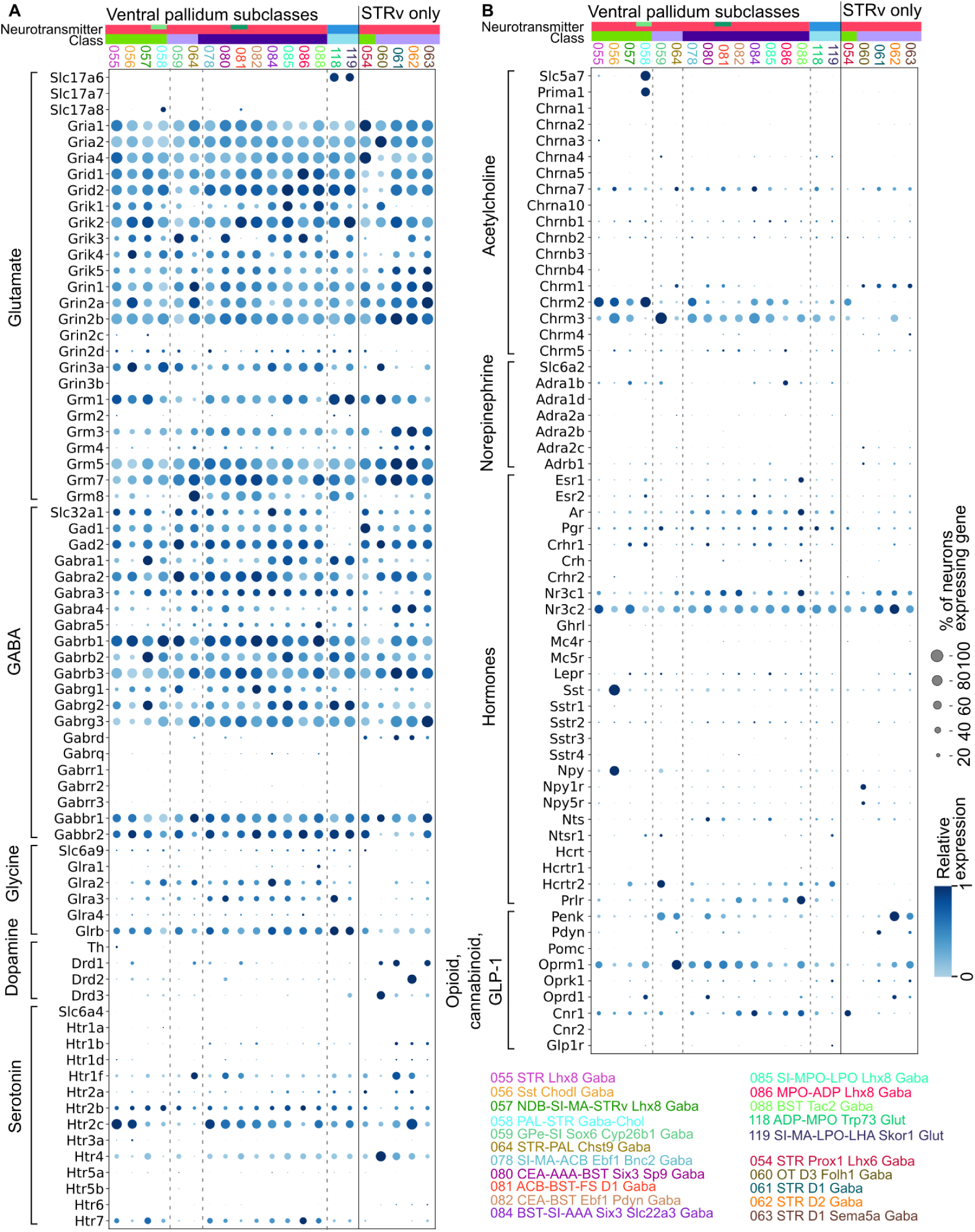
Expression of key neuronal signaling genes. (A) For each neuronal subclass, the fraction of neurons expressing (size) and normalized mean expression (color) of marker genes for the neurotransmitters glutamate, GABA, glycine, dopamine, and serotonin. Plotted for neurons from VP dissections in mouse and rat. We only plotted genes that were detected in our dataset. (B) As in (A), for acetylcholine, norepinephrine, selected hormones, opioids, cannabinoids, and GLP-1. We only plotted genes that were detected in our dataset.

**Figure S7.**
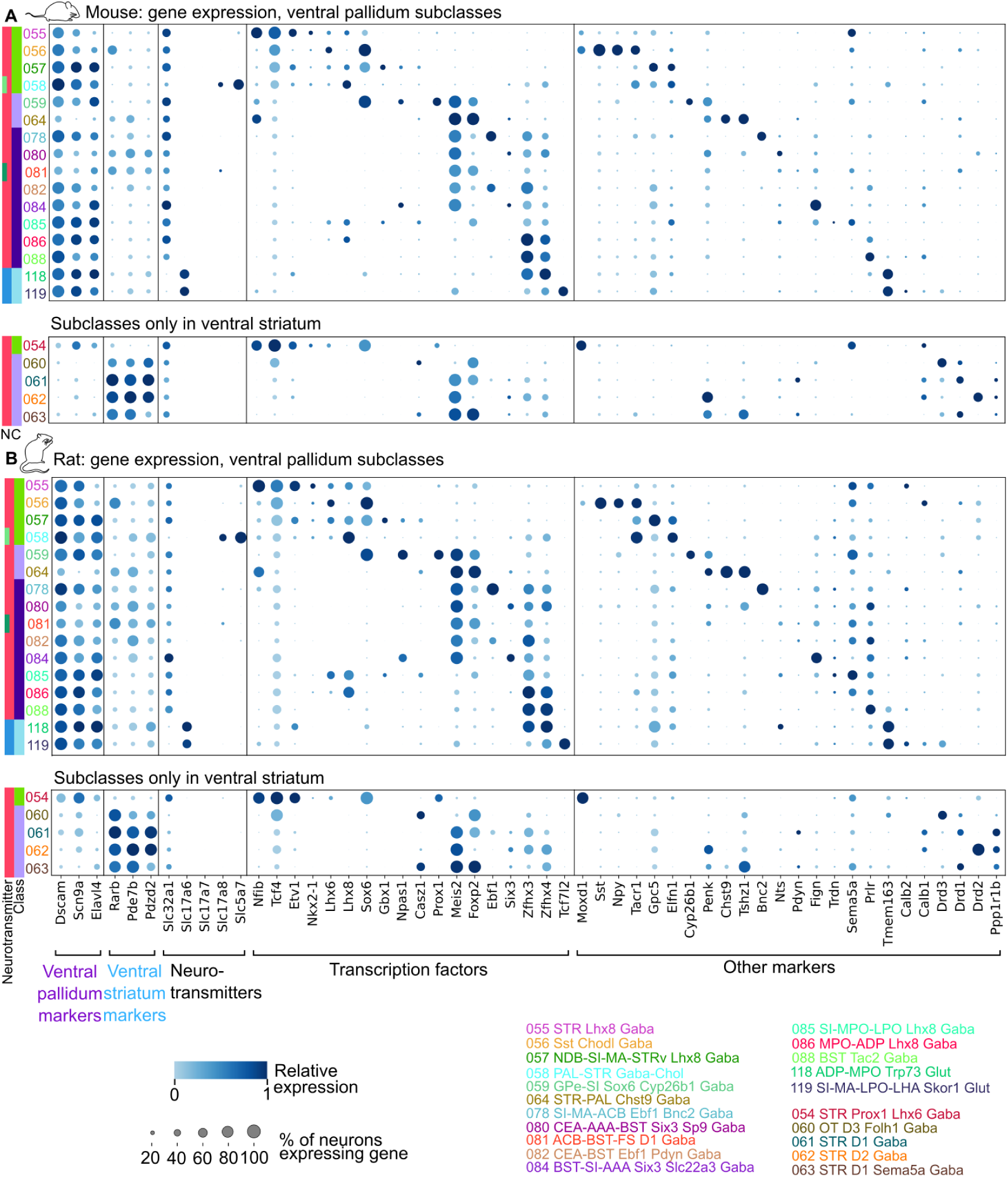
Comparison of VP expression patterns in mouse and rat. (A) For each neuronal subclass, the fraction of neurons expressing (size) and normalized mean expression (color) of marker genes of interest. Plotted for the neurons from VP dissections in mouse. (B) As in (A), for VP dissections in rat.

